# A curve fitting method for analysing starch granule size distributions in cereals

**DOI:** 10.1101/2024.10.03.616408

**Authors:** Rose McNelly, Amy Briffa, Georgia Yiasoumi, Cristobal Uauy, Ryo Matsushima, David Seung

**Affiliations:** John Innes Centre, Norwich Research Park, Norwich, NR4 7UH, UK; Institute of Plant Science and Resources, Okayama University, Kurashiki, 710-0046, Japan

## Abstract

**Background and Objectives:** The size distribution of starch granules is an important factor determining functional and nutritional properties of starch. However, a simple, standardised method for their analysis is lacking. Here, we developed an approach for estimating granule size parameters using a Python script that fits curves to volumetric granule size distributions generated using a Coulter counter.

**Findings:** The bimodal size distribution of starch from most wheat and barley cultivars could be best described with a mixed distribution curve. A log-normal distribution was fitted to the small B-type granules, and a normal distribution was fitted to the large A-type granules, allowing estimation of their relative abundance and size parameters, despite their overlapping size distributions. However, the optimal fitting is altered in wheat mutants with large perturbations in B-type granule content. In maize and rice, which have unimodal granule size distributions, size parameters were calculated by fitting a single normal distribution.

**Conclusions:** Curve fitting is an effective approach for estimating starch granule size parameters in diverse cereals, particularly the Triticeae with A- and B-type granules.

**Significance and novelty:** We provide new tools and guidelines for the quantitative analysis of granule size in cereals.

## 1 INTRODUCTION

The size distribution of starch granules is characteristic of the botanical source of the starch. In cereal endosperms, there is extensive natural variation in starch granule size distributions (Jane et al. 1994; Matsushima et al. 2013). Perhaps the most striking is the bimodal granule size distribution in grains of the Triticeae (wheat, barley and rye): where large A-type granules form during early grain development, and smaller B-type granules form in later grain development (Bechtel et al. 1990; Kamble et al. 2023). In wheat, B-type granules make up >90% of granules by number, but <30% by volume (Kamble et al. 2023; Lindeboom et al. 2004). Other cereals produce “simple” granules that grow from a single initiation and are relatively uniform in size (e.g., maize); or “compound” starch granules that develop when multiple granules initiate per amyloplast and grow against each other to form a larger, tessellated structure (e.g., rice) (Jane et al. 1994; Kawagoe 2013; Matsushima et al. 2015). Compound granules tend to fall apart during starch extraction, producing granules that are small, polygonal and relatively uniform in size. Some species, such as oats, have a more complex granule size distribution, containing both simple and compound-type granules (Saccomanno et al. 2017).

The size of starch granules has a major effect on the physicochemical and nutritional quality of starch (Li et al. 2021; Lindeboom et al. 2004). For example, the relative amount of B-type starch granules in wheat affects bread-and pasta-making quality (Park et al. 2009; Saccomanno et al. 2022; Soh et al. 2006), and in barley affects malting quality (Langenaeken et al. 2019). Larger granules are more resistant to digestion in the native state, as there is proportionally less surface area available for enzymatic attack (Dhital et al. 2010). Thus, methods to describe and quantify the size distribution of starch granules are important not only for the study of starch granule biosynthesis, but also for assessing starch quality for a broad range of applications.

Despite the importance of granule size distributions, there is currently no standardised method in grain research and industry to quantify it. However, several approaches have been used in the literature. A popular method is to measure granule size using images captured by light or electron microscopy (Jane et al. 1994). This requires relatively common instrumentation, and can quickly generate an estimate of the average granule size. However, it is labour intensive to generate high-quality size distributions using microscopy as many granules need to be quantified. Further, three-dimensional volume is difficult to determine from two-dimensional images, so the method generates granule diameter vs. number plots rather than a volumetric granule size distribution. Therefore, using a particle size analyser offers a far superior method to quantify granule size distributions in terms of accuracy, efficiency and reproducibility as they typically allow sizing of 10,000-100,000 particles in minutes. Such analysers include Coulter counters and light scattering particle size analysers. For accurate quantification of size, the Coulter counter is advantageous in that it directly measures the volume of granules using the electrochemical “Coulter principle”, and is relatively insensitive to granule shape (Lines 1981; Naito et al. 1998). In contrast, particle size estimates based on optical light/laser scattering are sensitive to particle shape (Barreiros et al. 2006; Naito et al. 1998), so differences in starch granule shape can distort the true measurement of volume. Regardless of the instrument of choice, both can provide volumetric granule size distributions, which must be further analysed to derive parameters of interest for statistical comparison between different samples (genotypes, treatments, developmental timepoints, etc).

Here, we developed a simple method for quantifying starch granule size distributions in various cereal grains, employing a Coulter counter and subsequent data analysis with a flexible Python script. We focused on the Coulter counter because it is more accessible and affordable than a light scattering device, and holds a significant advantage in that it directly measures volume with relative insensitivity to particle shape. The open-access Python script is provided online with documentation. We hope that the guidelines and resources in this article will lead to the development of more standardised methods for analysing starch granule size.

## 2 MATERIALS AND METHODS

### 2.1 Plant materials and growth

Seeds of the different wheat cultivars are from the John Innes Centre (JIC) Germplasm Resources Unit under the ‘Hexaploid wheat pangenome’ collection (accession codes PANG0001-PANG0016). The additional wheat cultivars Baj and Chinese Spring, barley cultivars Golden Promise, Nubet and Shikoku Hadaka, and *Aegilops tauschii* (accession AL8/78) were obtained from John Innes Centre Germplasm Resource Unit. Barley cultivar ‘Haruna Nijo’ was provided from NBRP-Barley (http://earth.nig.ac.jp/~dclust/cgi-bin/index.cgi?lang=en). Rice grains (Basmati) were purchased from a local market. Native maize starch was obtained from Merck (S4126).

All wheat cultivars were grown to provide seeds for starch extraction. These were grown in soil (John Innes cereal mix—65% peat, 25% loam, 10% grit, 3 kg/m^3^ dolomitic limestone, 1.3 kg/m^3^ pg mix, and 3 kg/m^3^ osmocote exact) in climate-controlled glasshouses. The glasshouses provided a minimum 16 h of light at 20 °C and 16 °C during the dark; and continuous relative humidity of 60%. For all other samples, starch was directly extracted from the obtained seeds.

### 2.2 Starch extraction

Seeds (2-3 seeds per sample) were soaked overnight in 0.5 M NaCl (0.5 mL), then homogenised for 1 minute in a ball mill (set to 25 Hz) with 3 mm tungsten carbide beads. The homogenate was filtered through a Pluriselect pluriStrainer Mini with 70 μm nylon mesh. The filter was washed with an additional 0.5 mL of ddH_2_O. Starch was pelleted at 3000*g* for 5 min. The pellet was washed three times in 1 mL 90% (v/v) buffered Percoll solution (prepared in 50 mM Tris-HCl, pH 8), spinning at 2500*g* for 5 min after each resuspension. The pellet was then washed three times in 1 mL of wash buffer (50 mM Tris-HCl, pH 6.8; 10 mM EDTA; 4% SDS; and 10 mM DTT), spinning at 5000*g* for 2 min after each resuspension. Finally, the pellet was washed three times in 1 mL of distilled water, spinning at 5000*g* for 1 min after each resuspension.

### 2.3 Coulter Counter analysis

Purified starch granules were resuspended in Isoton II electrolyte (Beckman Coulter) and analysed on a Multisizer 4e Coulter counter (Beckman Coulter), fitted with a 70 μm aperture and 100 mL beaker. The amount of starch measured was optimised such that it did not exceed the 10% concentration bar, to ensure that co-incidences (two particles passing through the aperture at once) were minimised. The Coulter counter was set to measure 50,000 particles (starch granules) per sample and all measurements were performed using logarithmic bin spacing. The data were exported to Comma Separated Value (CSV) files containing the relative volume measured in each bin. For consistency with previous literature in the field, the Coulter Counter was set to output bins measured in units of particle diameter. As the Coulter counter directly measures particle volume, this involves a spherical particle assumption. For naturally spherical granules, such as B-type wheat granules, this is a good approximation. For those with more complex geometries, however, such as the lenticular A-type wheat granules or the polyhedral granules of rice, we note that this is an effective diameter for a spherical granule of equivalent volume.

### 2.4 Data input into curve-fitting script

The sections below outline the development and operation of the script, which were written in Python 3.11.5. The CSV output files from the Coulter counter were directly input into the script to generate a volumetric granule size distribution (Figure 1a). To prepare data for curve fitting, the script normalises the data by dividing the relative volume in each bin by the bin width, to account for the logarithmic bin spacing during data collection. The curve fitting below is performed on these normalised data (Figure 1b). If the data were acquired with linear (even) bin spacing, this bin width normalisation will not change the distribution.

**Figure 1:**
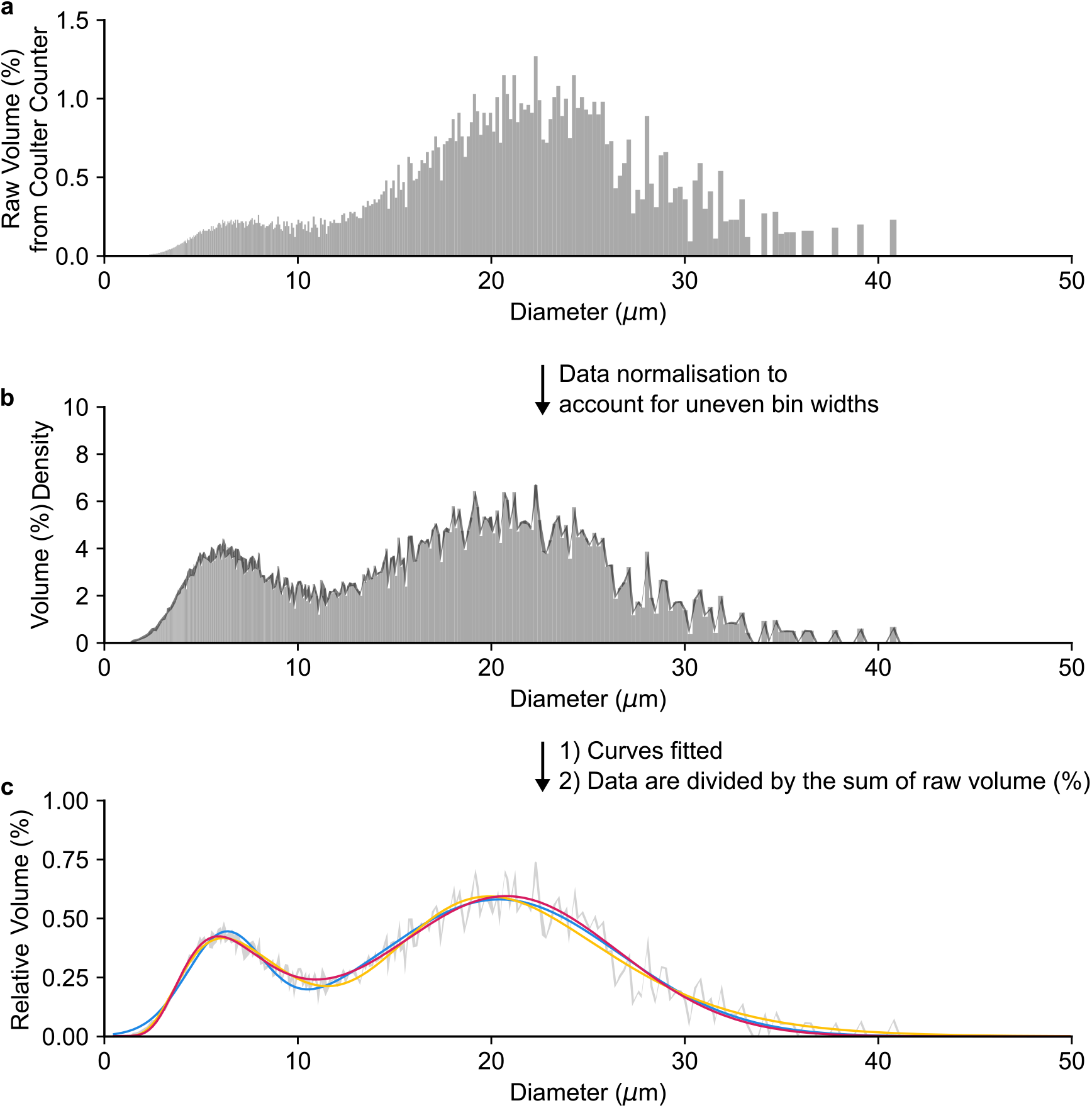
Transformation of Coulter counter data for accurate fittings. **(a)** Raw volumetric starch granule size data from wheat cultivar Claire, quantified using a Coulter counter with logarithmic bin spacing. Here it is plotted over a linear scale as a histogram. Logarithmic bin spacing means that bins at smaller diameters are narrower than those at larger diameters, so peak heights must be normalised to account for this. **(b)** Volumetric granule size distribution data from (a) after normalisation for logarithmic bin spacing. A line (black) has been overlaid with the volume (%) density plotted against the midpoint of the bin, this is the curve which the fitting is conducted against. **(c)** After curve fitting, the data (grey) and the fitted curves are divided by the sum of the raw volume (%) so that the total height of the data sums to 100%. This is a simple stretch and results in no changes to the overall fitting or calculated parameters. This ensures the units of the y-axis is in line with expectations from the output of the Coulter counter. Fitted curves are in blue (normal-normal), yellow (log-normal-log-normal) or pink (log-normal-normal).

A generic data input format is provided together with script documentation, such that data from any Coulter counter model can be used for the analyses.

### 2.5 Defining fittings to bimodal curves

To perform fits with a normal-normal (N-N) function:

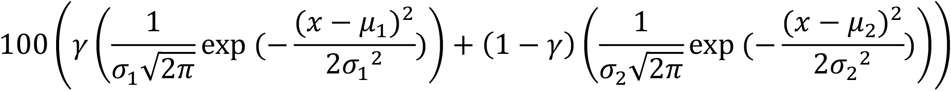

the initial parameters were: γ = 0.2, μ_1_ =5,σ_1_ = 2, μ_2_ = 20,σ_2_ =5. The value of γ was constrained to 0 ≤ γ ≤ 1, whilst all others were constrained between 0 and ∞.

To perform fits with a log-normal–log-normal (L-L) function:

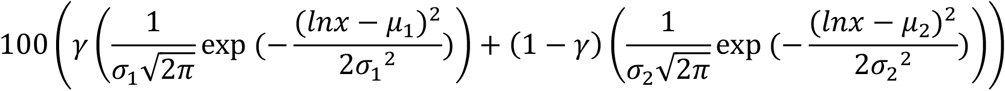

The initial parameters were: γ = 0.5, μ_1_ = 2,σ_1_ = 0.5, μ_2_ = 3,σ_2_ = 0.2. The value of γ was constrained to 0 ≤ γ ≤ 1, whilst all others were constrained between 0 and ∞.

To perform fits with a log-normal-normal (L-N) function:

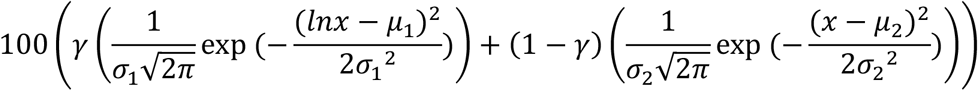

the initial parameters were: γ = 0.2, μ_1_ = 1.5,σ_1_ = 0.5, μ_2_ = 20,σ_2_ =5. The value of γ was constrained to 0 ≤ γ ≤ 1, whilst all others were constrained between 0 and ∞.

The initial parameters were selected as they produced curves which closely followed a typical normalised coulter counter trace produced by wheat starch (Figure 1b). In all the above fits, 1 corresponds to the B-type granule distribution and 2 corresponds to the A-type granule distribution.

### 2.6 Defining fittings to unimodal curves

To perform fits with a normal function:

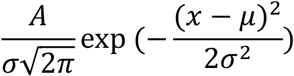

The initial parameters were: *A* = 100, μ =5,σ = 2, and all parameters were constrained between 0 and ∞.

To perform fits with a log-normal function:

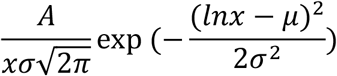

The initial parameters were: *A* = 200, μ = 2,σ = 1, and all parameters were constrained between 0 and ∞.

The initial parameters were selected as they produced curves which followed a typical Coulter counter trace produced by maize and rice starch.

### 2.7 Optimisation of fittings and calculation of parameters

For all fittings, the optimise.curve_fit function from the scipy 1.11.1 package (Virtanen et al. 2020) was used to fit optimised distributions, using the previously stated initial values as the starting points. The mean and variance of all mathematical curves as quoted throughout this work are calculated given the standard calculations for normal and log-normal curves. We note that these values do not directly equate to the mean and standard deviation of the particle-size distribution, but nevertheless, provide an excellent proxy for comparing size parameters between different samples. For bimodal fittings, the B-type granule content (%) is given by the value of 100 ∗ γ. Subsequently, the A-type granule content (%) is calculated as 100 – B-type granule content (%).

For each fitting, the standard error of regression (S) is provided as a goodness of fit test, which was calculated as:

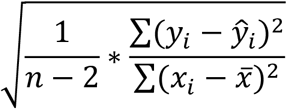

where *n* is the total number of data points, *y*_*i*_ the measured *y* value, *ŷ*_*i*_ the *y* value calculated from the fitting, *x*_*i*_ the measured *x* value and 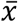 the mean *x* value. In addition, we calculated individual parameter uncertainties as the square root of the diagonal of the covariance matrix of errors produced by optimise.curve_fit. Total fit uncertainty was calculated as the sum of all parameter uncertainties.

### 2.8 Plotting fits and script outputs

The scripts produce multiple outputs to allow efficient data analysis.

Firstly, it generates a series of pdf files containing graphs of the fittings for each of the samples. In these graphs, the data and fitted curves have been divided by the sum of the raw volume (%), which is a simple stretch and results in no changes to the overall fitting or calculated parameters (Figure 1c). This brings the *y*-axis more in line with the scale from the raw output of the Coulter counter and previous literature.

Secondly, it produces Excel spreadsheets: ‘Starch_parameters_from_fittings’ which gives the mean and variance of the fitted volumetric granule size distributions for all optimised fittings, and for the bimodal script gives A- and B-granule contents (Table 1); ‘Mathematical_values_to_reproduce_fittings’ contains all fitting parameters from the optimised fitting, along with uncertainties for each fitting parameter, to allow users to re-plot the mathematical curves if required (Table 2).

**Table 1.** Example of the starch parameter output from the bimodal fitting script for the L-N fitting to starch from wheat cultivar Claire. Diameters and variances are quoted in the same units as the bin-specifications of the original data file, in this case μm.

| Sample | Mean of the B-type granule diameter-distribution | Mean of the A-type granule diameter-distribution | B-type granule content (%) | A-type granule content (%) | Variance of B-type granule diameter-distribution | Variance of A-type granule diameter-distribution | Fit Uncertainty | Standard error of regression |
| --- | --- | --- | --- | --- | --- | --- | --- | --- |
| Claire | 7.246 | 20.813 | 22.241 | 77.759 | 8.529 | 33.214 | 0.219 | 0.002 |

**Table 2.** Example of the mathematical output from the bimodal fitting script for the L-N fitting to starch from wheat cultivar Claire.

| Sample | $\gamma$ | $\gamma$ uncert | $\mu_1$ | $\mu_1$ uncert | $\sigma_1$ | $\sigma_1$ uncert | $\mu_2$ | $\mu_2$ uncert | $\sigma_2$ | $\sigma_2$ uncert |
| --- | --- | --- | --- | --- | --- | --- | --- | --- | --- | --- |
| Claire | 0.222 | 0.007 | 1.905 | 0.013 | 0.388 | 0.009 | 20.813 | 0.091 | 5.763 | 0.098 |

Finally, the script produces a plot of S and total fit uncertainty values for all models for all samples. This assists the user in selecting the best overall model for curve fitting within their dataset, as the model that produces the lowest values for most samples can be easily visualised.

Error catching is incorporated to produce lists of samples where 1) the script completely failed to produce fittings (“Samples_where_the_script_failed”), suggesting an issue with the format of the input data, or 2) where the fitting could not be sufficiently optimised (“Samples_which_fitted_successfully_and_samples_which_failed”) suggesting that none of the models provide an optimal fit for the data.

### 2.9 Code availability

The scripts and instructions for use are freely available online at https://github.com/DavidSeungLab/Coulter-Counter-Data-Analysis. An example of how data should be organised to use the script is also included.

## 3. RESULTS

### 3.1 A mixed log-normal/normal distribution fits starch granule size distributions of wheat

We developed a Python script to estimate A- and B-type granule contents of wheat starch by calculating the relative area under the A- and B-type granule peaks in Coulter counter data. These two peaks are not completely resolved and have substantial overlap, thus requiring curve fitting to separate them for calculating their relative areas (Figure 1). In this regard, a previous study developed a script for fitting mixed distributions to wheat starch granule size distributions from a laser scattering instrument, where the curve was a mixture of three log-normal distributions (Tanaka et al. 2017). Two of the three distributions correspond to A- and B-type granules, and the third corresponds to so called “C-type” granules, which is defined as a subpopulation of granules that is smaller than the B-type granules. However, when designing our script, we opted to fit bimodal distributions for the analysis of wheat starch, excluding consideration of C-type granules. Firstly, only a few studies report the presence of C-type granules (Bechtel et al. 1990; Wilson et al. 2006), and in our own analysis, we only observed peaks corresponding to A- and B-type granules (Figure 1). Secondly, unlike the A- and B-type granules that arise from two defined waves of granule initiation, the biological origin of C-type granules remains unexplained. Finally, the contribution of C-type granules to starch quality is unclear, since they have an almost negligible contribution to the total granule volume due to their small size and low abundance. Our script fits three different mixed distribution curves to the bimodal starch granule size distribution, including normal distributions to both A- and B-type granule peaks (henceforth called N-N), log-normal distributions to both peaks (henceforth called L-L), and a log-normal distribution to the B-type granule peak and a normal distribution to the A-type granule peak (henceforth called L-N)(Figure 2). The reason why we considered fitting a log-normal distribution to the B-type granule peak is because the lower end of the peak comes close to the y-axis and there can be no granules with a negative diameter value. Full description of the curve fitting is described in Sections 2.4-2.9.

**Figure 2:**
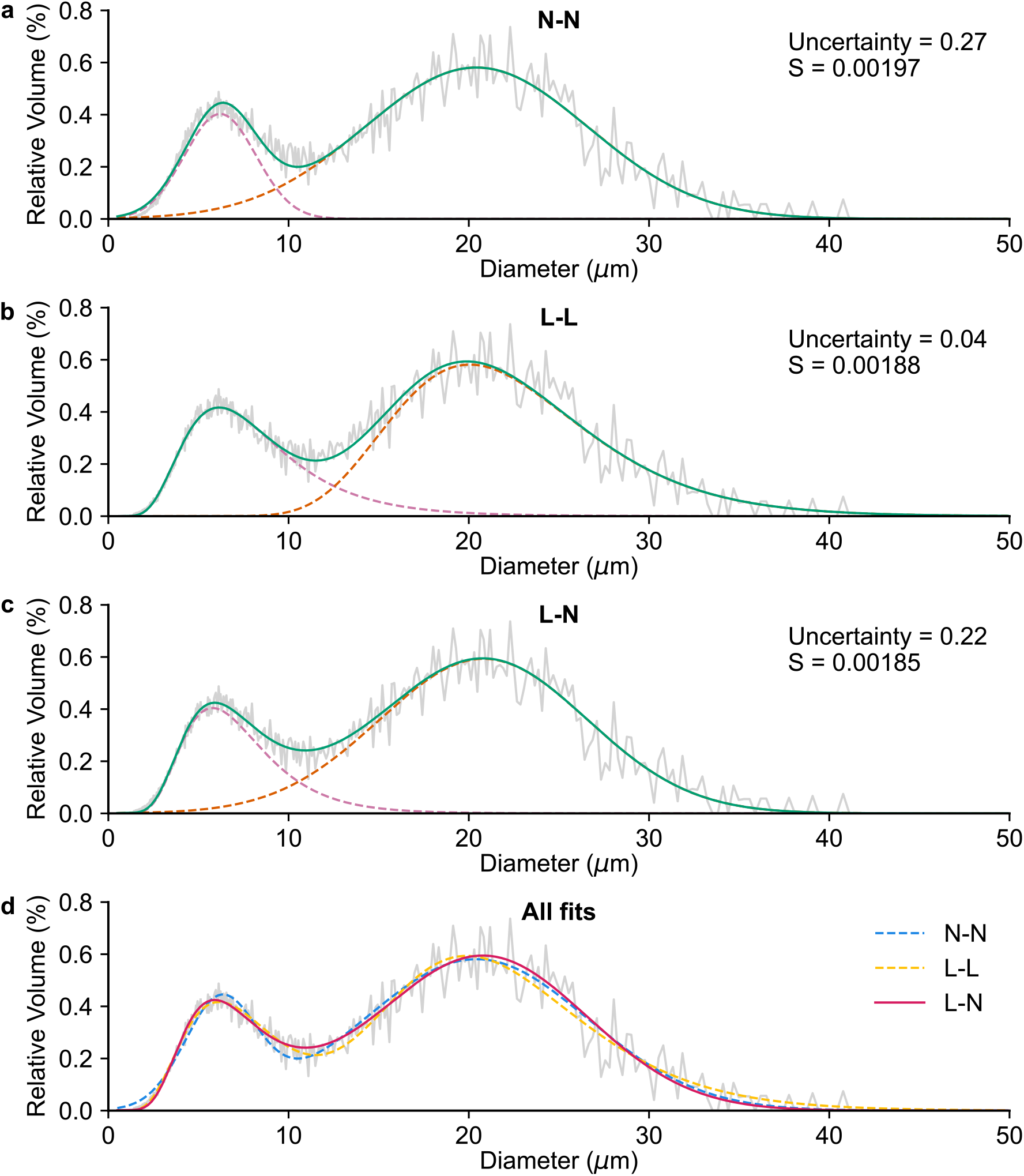
Three different bimodal curve fittings to the volumetric distribution of wheat starch. The normalised volumetric size distribution was determined using a Coulter counter on starch from wheat cultivar Claire (grey). **(a-c)** We fit two normal (N-N) (a), two log-normal (L-L) (b), or a log-normal and a normal (L-N) (c) distributions to the measured size distribution. The orange dashed curve represents the fitted A-type granule distribution, the pale pink dashed curve represents the fitted B-type granule distribution, and the green curve is the sum of these two distributions. The fit uncertainty and standard error of regression (S) are reported in the top right. **(d)** All three fittings have been overlaid for comparison, with the fitting with the lowest S value as a solid line and the other fittings as dashed lines.

We tested the fitting of the three different curves on granule size distributions from wheat starches. Initially, we assessed the quality of fitting using the standard error of regression (S) and fit uncertainty values provided by the script. In a typical wheat starch sample from cultivar Claire, the uncertainty values were low for all fits. The S values were also low but lowest for the L-N fitting, indicating it was mathematically the best fit (Figure 2a-c). Visual assessment of the fitted curves showed that the N-N curve deviated substantially from the actual distribution at the lower size range (between 0-3 μm) (Figure 2d). In contrast, the L-L curve deviated from the distribution at the upper size range (between 32-40 μm). However, a major difference between the L-L and the N-N fitting was that in the L-L fitting, we observed a long “tail” on the upper end of the B-type granule peak (between 12-20 μm), which was accommodated by the shape of the log-normal A-type granule peak. This tail implies that a substantial population of B-type granules are between 12-20 μm, which is almost never observed using microscopy. The L-N curve fitting minimised these distortions. In the L-N fitting, the lower end of the B-type peak and the upper end of the A-type granule peak closely followed the actual granule distribution trace, and fixing the A-type granule peak to a normal distribution reduced the “tail” of the log-normal B-type granule peak. Thus, from both our mathematical and visual assessments, the L-N was the optimum fit.

To test the how these curve fittings might vary with genotypic variation in granule size distributions in wheat, we analysed fittings for 65 different Coulter counter traces representing 17 diverse wheat cultivars from the 10+ wheat genomes project (Walkowiak et al. 2020), the genome reference Chinese Spring, and the CIMMYT cultivar Baj (Figure 3; Table S1, S2). We saw large differences in the size distribution traces, particularly in the position of the peaks, and thus the mean granule diameters (Figure 3, S1). Uncertainty values were low (< 0.75) for all three fits. Comparison of S values showed that the majority of samples (60%) produced lowest S values with the L-N fitting, while 40% had lowest S value for the L-L fitting (Figure 3e). The N-N distribution never produced the lowest S value.

**Figure 3:**
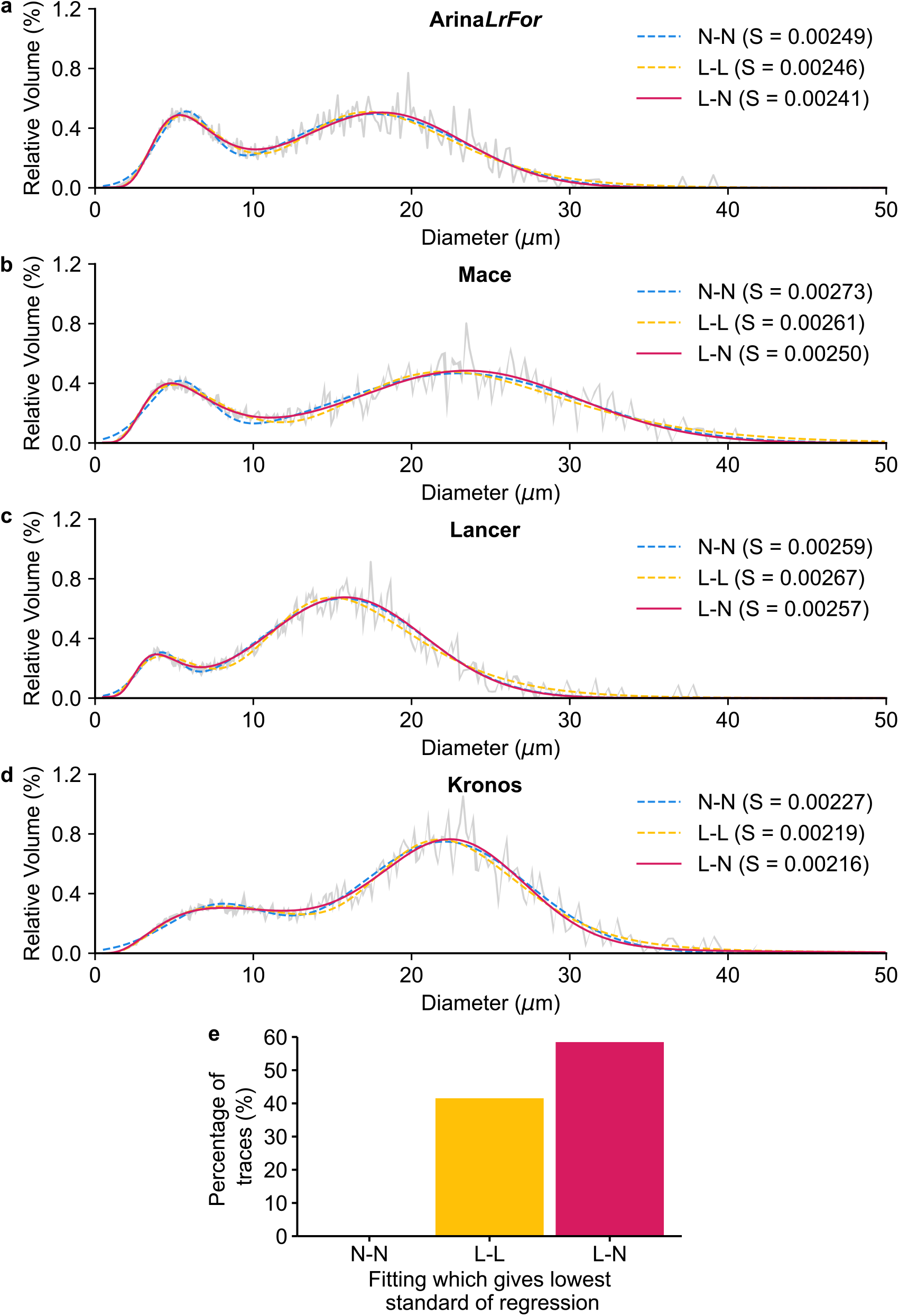
Comparison of the three different fittings with starch from different wheat cultivars. **(a-d)** The normalised volumetric size distribution was determined using a Coulter counter on starch from bread wheat cultivars Arina*LrFor* (a), Mace (b), Lancer (c) and durum wheat Kronos (d) (all grey). We fit two normal (N-N, blue), two log-normal (L-L, yellow) or a log-normal and normal distributions (L-N, pink) to the measured size distribution, with the fit with the lowest standard error of regression (S) value represented as a solid line and other fits as dashed lines. **(e)** Sixty-five (*n)* coulter counter traces from 17 wheat varieties were run through the fitting script, and the fitting with the lowest S value was calculated and is shown as a percentage.

From these fittings, we derived the mean of the fitted A-type granule distribution curve, mean of the fitted B-type granule distribution curve, and B-type granule content by volume (the integral of the B-type granule peak as a percentage of the total integral of both curves). To compare values derived from the different fittings, values from the L-N fitting were plotted against those from the N-N and double L-L fittings (Figure 4). Generally, the values from the three different fittings for each sample were consistent with each other in rank, such that samples having high values under one fit were also high with the other fits. Thus, the Pearson correlation coefficients between values were all greater than 0.85. However, the absolute values obtained for each parameter varied substantially between the different fittings. The mean A-type granule diameter-distribution was similar between the L-N and N-N fittings. The L-L fitting, however, resulted in higher diameters for both A- and B-type granules. This is likely a result of the long tail observed on the B-type granule peak under the L-L fitting, which also shifts the position of the A-type granule peak. The L-L fitting also produced the highest B-type granule content values, which is again likely an overestimate produced by the long tail.

**Figure 4:**
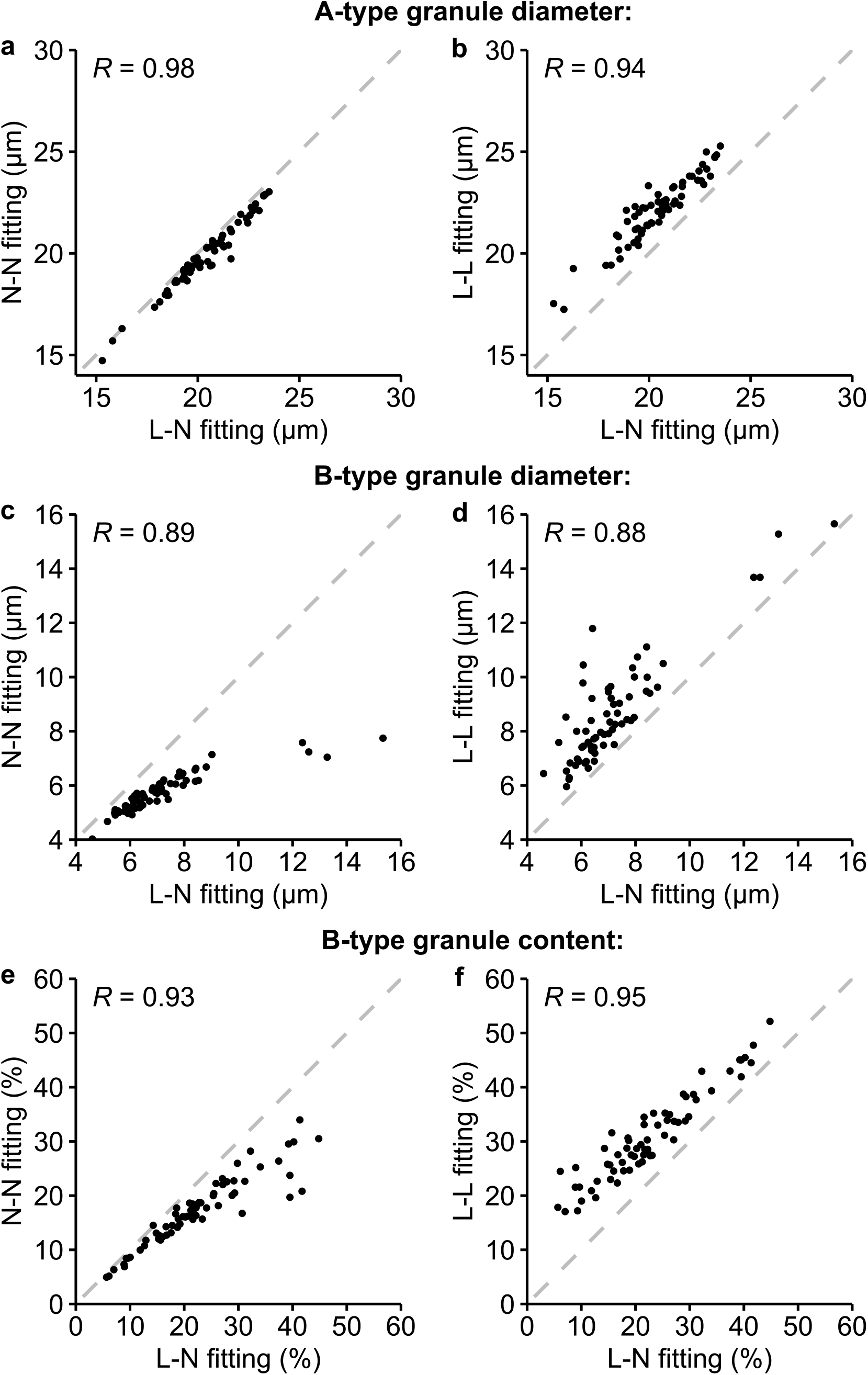
Comparison of parameters derived from curve fittings. A-type granule diameter-distribution mean (a-b), B-type granule diameter-distribution mean (c-d) and B-type granule content (e-f) were derived from the normal-normal N-N (a, c, e), log-normal-log-normal L-L (b, d, f) fittings, and plotted against those derived from the log-normal-normal L-N fittings (a-f). Individual data points are shown as black dots (*n* = 65), with the Pearson correlation coe[icient shown in the top left. A dashed grey line of *y*=*x* is overlaid on each plot to show the expected relationship if the fittings gave the same values.

It therefore appears that for a broad range of wheat cultivars, the L-N fitting provided a reliable fit that is consistent with experimental observations. We next tested whether the fitting would also work on extreme, induced variation in B-type granule content. We analysed starch from durum wheat mutants deficient in the plastidial *Phosphorylase 1* (*phs1*) and *Myosin-resembling Chloroplast Protein* (*mrc*), which have low and high B-type granule contents, respectively (Figure 5; Table S3)(Chen et al. 2024; Kamble et al. 2023). Interestingly, the L-N fitting did not provide the best fit for either mutant. For *mrc*, the L-L consistently fitted best. For *phs1*, the best fit was more variable: the N-N fitting was best for 50% of the samples, followed by L-L and L-N. However, it should be noted that the S values were small for all fittings on both mutants, with the maximum S value observed being S=0.0022.

**Figure 5:**
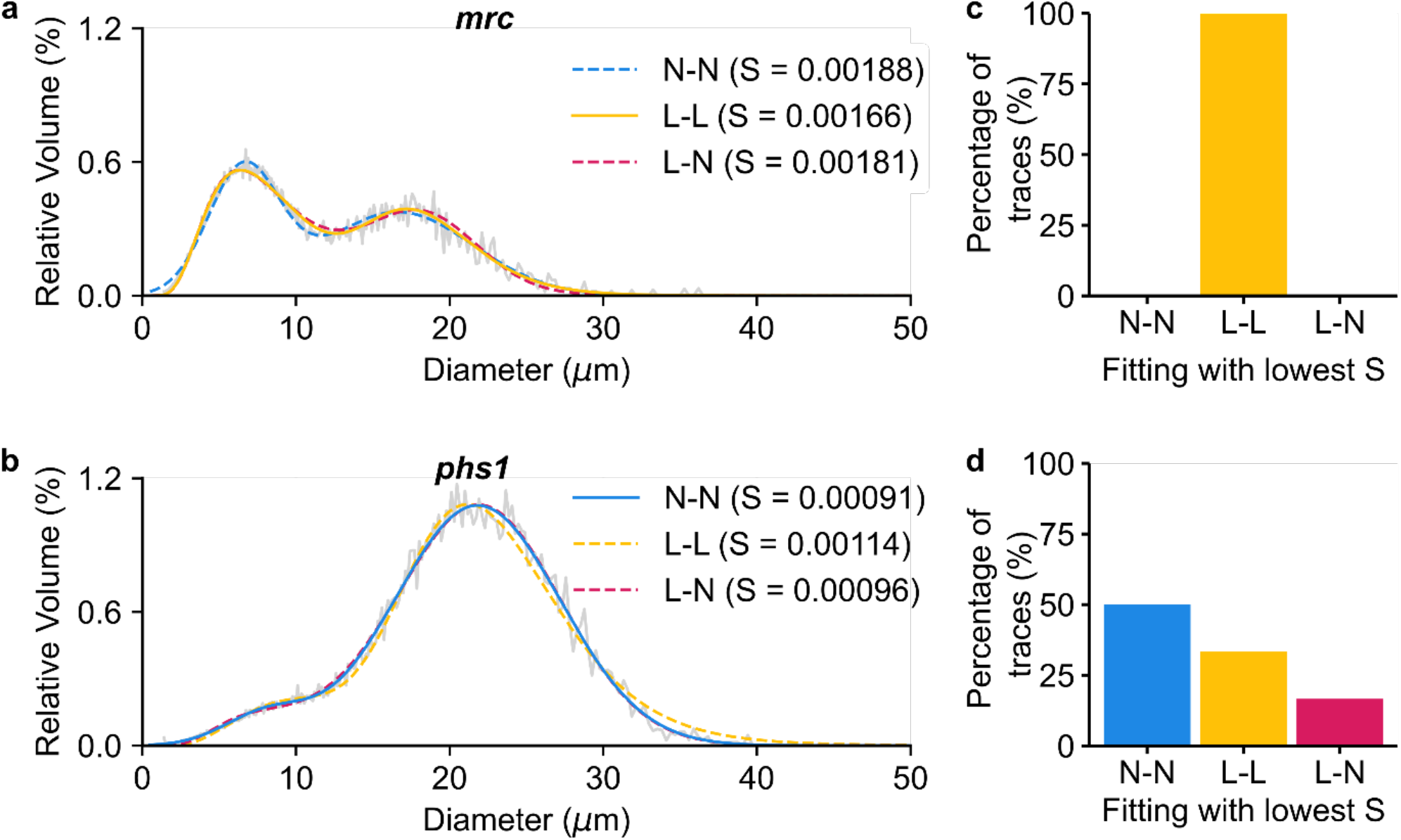
Comparison of the three different fittings with wheat mutants with altered B-type granule contents. **(a-b)** The normalised volumetric size distribution was determined using a Coulter counter on starch from *mrc* (a), *phs1* (b) (both grey). We fit two normal (N-N, blue), two log-normal (L-L, yellow) or a log-normal and normal (L-N, pink) distributions to the measured size distribution, with the fit with the lowest standard error of regression (S) value represented as a solid line and other fits as dashed lines. **(c-d)** The fittings with the lowest S value for *mrc* (c, *n* = 9) and *phs1* (d, *n* = 6) were calculated and are shown as percentages.

### 3.2 The L-N distribution fits starches from diZerent Triticeae species

Given that the L-N model fit best to most natural wheat varieties, we assessed whether this distribution could also fit starch granule size distributions from other Triticeae species. We tested starch from four different barley cultivars (Golden promise, Nubet, Haruna Nijo and Shikoku Hadaka) and the wild wheat relative, *Aegilops tauschii*. The L-N fitting produced the most optimal fitting for all these samples (Figure 6, S2; Table S4).

**Figure 6:**
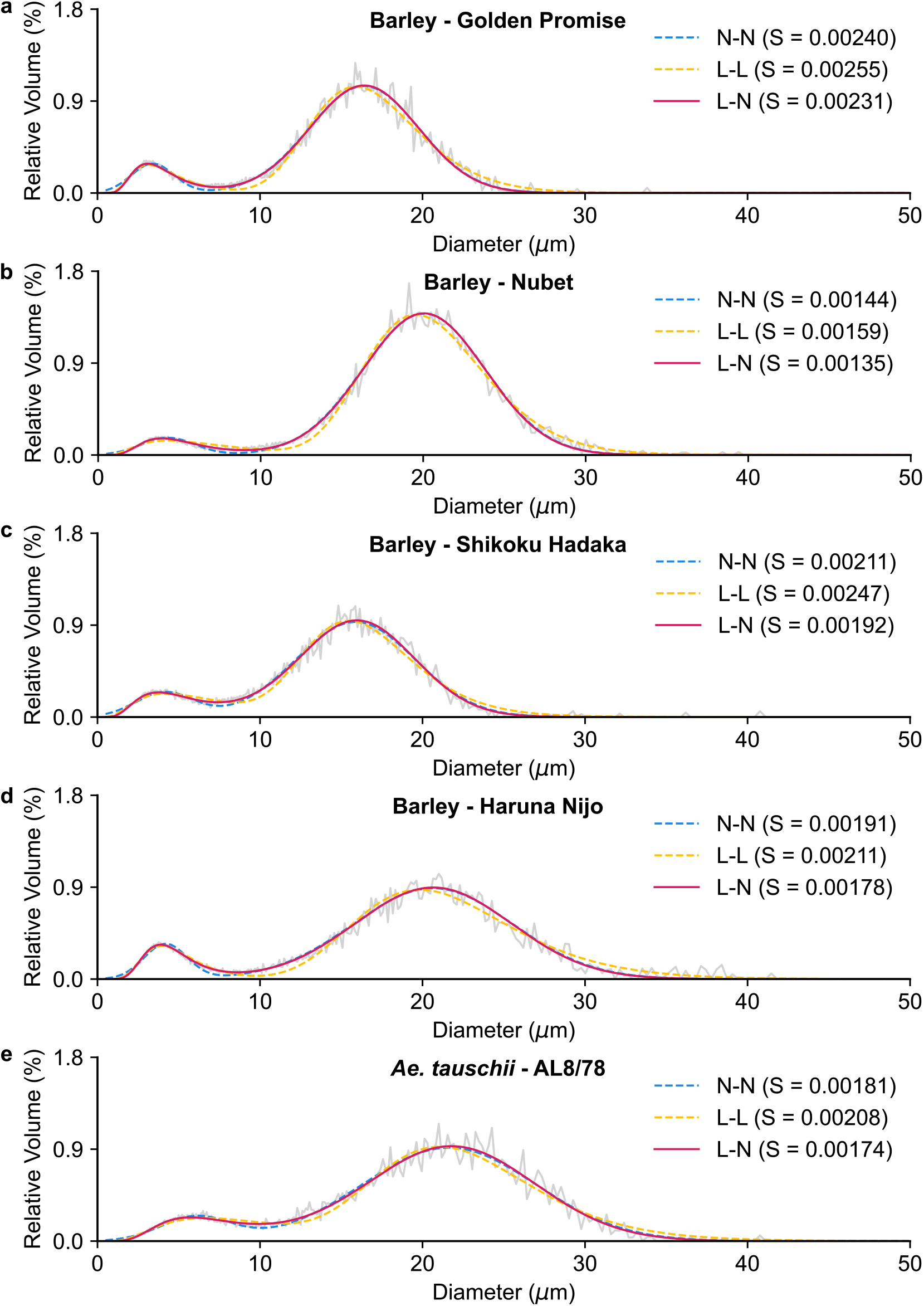
Comparison of the three different fittings with different Triticeae starches. The normalised volumetric size distribution was determined using a Coulter counter on starch from barley cultivars Golden promise (a), Nubet (b), Haruna Nijo (c) and Shikoku Hadaka (d), and the *Ae. tauschii* genome reference AL8/78 (e) (all grey). We fit two normal (N-N, blue), two log-normal (L-L, yellow) or a log-normal and normal (L-N, pink) distributions to the measured size distribution, with the fit with the lowest standard error of regression (S) value represented as a solid line and other fits as dashed lines.

### 3.3 Adaptation of the script to analyse starch with a unimodal distribution

Since the occurrence of A- and B-type granules is restricted to the Triticeae, we created a version of the script for fitting a unimodal granule size distribution. This script performs bin width normalisation as described for the bimodal script, and fits both normal and log-normal distributions to the data. We tested the script on maize and rice starch, and for both the normal distribution fit better than the log-normal distribution, producing a visually better fit and low fit uncertainty and S values (Figure 7, Table S5). However, in rice, we observed the presence of some large particles (>8 μm) that extended beyond the fitted normal distribution (Figure 7).

**Figure 7:**
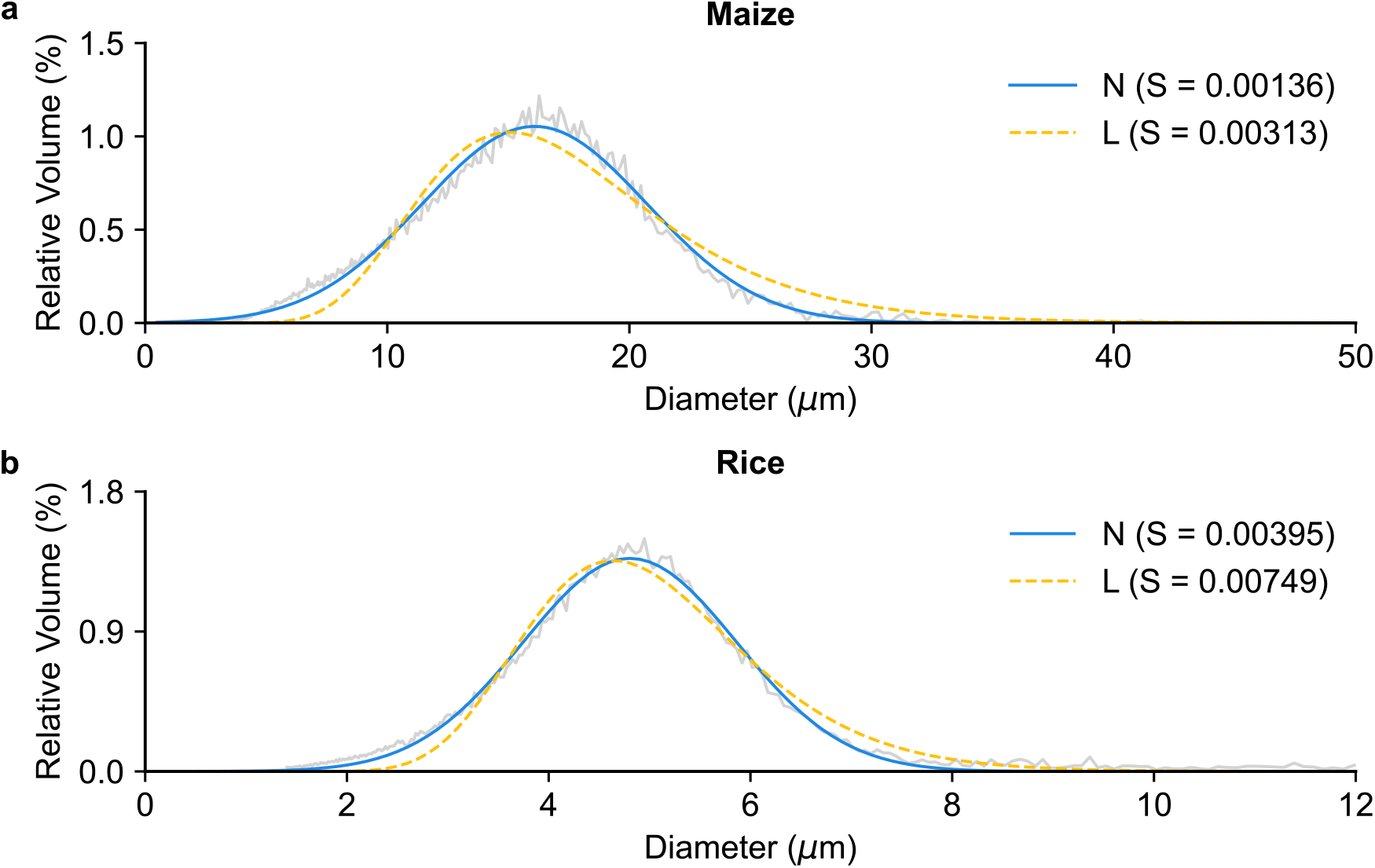
Comparison of the two different fittings for starch with a unimodal size distribution. The normalised volumetric size distribution was determined using a Coulter counter on starch from maize (a) and rice (b) (both grey), note that the x scale has been adjusted in (b) as the rice starch is smaller. We fit a normal (N, blue) and a log-normal (L, yellow) curve to each measured size distribution, and the curve with the lower standard error of regression (S) value is represented with a solid line, whereas the other is shown with a dashed line.

## 4 DISCUSSION

Our method for quantifying starch granule size distributions combines Coulter counter measurements with a simple analytical script. This allows size distributions obtained from large populations of starch granules to be summarised as size parameters. We demonstrated the applicability of this approach to different cereals, including various wheat and barley cultivars, *Aegilops*, maize and rice. We have summarised the granule size parameters obtained from these analyses in Table 3, showing differences in granule size parameters between wheat and barley cultivars. Small differences between varieties cannot be easily distinguished using microscopy, demonstrating the usefulness of our approach.

**Table 3.** Granule size parameters derived from the Coulter counter followed by curve-fitting analysis. Values are mean ± SEM from *n*=3-4 replicates, where each represents grains harvested from different plants. Where only one seed batch was available (*), value represents a single analysis of from starch preparation from pooled grains from separate plants.

| | Mean of the B-type granule diameter-distribution ($\mu\text{m}$) | Mean of the A-type granule diameter-distribution ($\mu\text{m}$) | B-type granule content (%) | A-type granule content (%) |
| --- | --- | --- | --- | --- |
| **Bimodal fittings (L-N)** |  |  |  |  |
| Wheat cultivars |  |  |  |  |
| Range: | (5.85 – 8.39) | (17.63 – 22.52) | (7.61 – 35.40) | (64.60 – 92.39) |
| ArinaLrFor | 6.23 ± 0.30 | 17.85 ± 0.98 | 27.67 ± 1.56 | 72.33 ± 1.56 |
| Baj | 5.94 ± 0.21 | 19.39 ± 0.05 | 7.61 ± 1.00 | 92.39 ± 1.00 |
| Cadenza | 6.18 ± 0.31 | 19.44 ± 0.44 | 16.24 ± 2.04 | 83.76 ± 2.04 |
| CDC-Landmark | 8.00 ± 0.28 | 20.61 ± 0.23 | 35.40 ± 3.15 | 64.60 ± 3.15 |
| CDC-Stanley | 7.62 ± 0.62 | 20.22 ± 0.38 | 28.36 ± 4.43 | 71.64 ± 4.43 |
| Chinese Spring | 7.35 ± 0.21 | 20.80 ± 0.81 | 20.03 ± 2.16 | 79.97 ± 2.16 |
| Claire | 6.47 ± 0.37 | 20.24 ± 0.62 | 24.59 ± 2.61 | 75.41 ± 2.61 |
| Jagger | 6.87 ± 0.41 | 20.33 ± 0.93 | 19.99 ± 1.40 | 80.01 ± 1.40 |
| Julius | 7.03 ± 0.21 | 22.52 ± 0.35 | 25.49 ± 1.57 | 74.51 ± 1.57 |
| Kronos | 6.95 ± 0.21 | 22.02 ± 0.06 | 25.84 ± 0.86 | 74.16 ± 0.86 |
| Lancer | 5.85 ± 0.82 | 17.63 ± 0.91 | 19.11 ± 4.79 | 80.89 ± 4.79 |
| Mace | 6.00 ± 0.42 | 20.93 ± 1.30 | 16.59 ± 4.79 | 83.41 ± 4.79 |
| Norin61 | 6.15 ± 0.24 | 18.84 ± 0.91 | 16.80 ± 2.07 | 83.20 ± 2.07 |
| Paragon | 6.58 ± 0.30 | 21.50 ± 0.74 | 18.47 ± 2.02 | 81.53 ± 2.02 |
| Robigus | 8.39 ± 0.31 | 21.90 ± 0.47 | 33.03 ± 5.74 | 66.97 ± 5.74 |
| SY-Mattis | 7.08 ± 0.39 | 21.10 ± 0.78 | 14.44 ± 1.78 | 85.56 ± 1.78 |
| Weebil | 6.33 ± 0.24 | 20.83 ± 0.16 | 17.36 ± 3.48 | 82.64 ± 3.48 |
| Barley cultivars |  |  |  |  |
| Range: | (4.02 – 5.46) | (15.98 – 20.25) | (5.82 – 15.24) | (84.76 – 94.18) |
| Golden Promise | 4.02 ± 0.07 | 16.33 ± 0.25 | 10.79 ± 0.75 | 89.21 ± 0.75 |
| Nubet* | 5.31 | 20.05 | 5.82 | 94.18 |
| Haruna Nijo | 4.57 ± 0.05 | 20.25 ± 0.63 | 15.24 ± 1.80 | 84.76 ± 1.80 |
| Shikoku Hadaka* | 5.46 | 15.98 | 13.34 | 86.66 |
| Other species |  |  |  |  |
| *Ae. tauschii* | 7.51 ± 0.56 | 21.87 ± 0.16 | 11.52 ± 0.63 | 88.48 ± 0.63 |
| **Unimodal fittings (N)** |  |  |  |  |
| | Mean of the granule diameter-distribution ($\mu\text{m}$) | | | |
| Maize | 16.07 ± 0.01 | | | |
| Rice | 4.80 ± 0.00 | | | |

To allow this approach to be easily applied to a range of different samples, we provided two different scripts for fitting unimodal and bimodal distributions, and each can fit normal and log-normal distributions. Even within the small subset of cereal species analysed here, we observed that different fittings worked best depending on species and genotype. For example, L-N mixed distributions fit the majority of wheat and barley cultivars, and the wild wheat relative *Ae. tauschii*. However, when we perturbed the B-type granule content by introducing mutations in wheat, the L-N no longer provided the best fit. Maize and rice starch with unimodal distributions could be fitted with a single normal distribution. The use of L-N distributions for describing Triticeae starch and the fact that starch from different species, cultivars and mutants fit different distributions, are major findings of our work.

Our script works by fitting known mathematical distributions to the experimentally determined granule size distribution. However, the biological reason why different fittings work better for different species is unknown. It is important to note that the shape of the size distribution is also influenced by technical factors. For example, although the Coulter counter measures the volume of each particle and is insensitive to shape, it assumes spherical particle size when it calculates a diameter value for the x-axis. A-type granules, however, are known to be lenticular. Outputting Coulter Counter data using bin-volumes (as opposed to diameter) and fitting accordingly would provide a more direct quantification of starch granule size properties. However, granule size is conventionally reported in the literature as diameter rather than volume, and the script as currently presented here has the advantage of allowing direct comparisons to the existing literature.

It is essential that the user selects the fitting that is optimal for their experimental data, and we provide two mathematical indicators (S and fit uncertainty) which can guide the selection. The S value, which assesses goodness of fit, should drive model selection. However, S values should only be used to compare the different bimodal fittings or different unimodal fittings and cannot be used to compare unimodal and bimodal fittings, due to the different number of parameters in the models. Visual inspection of the curves is essential to determine if bimodal or unimodal fitting is appropriate, and to assess the quality of the fitting. It is also important to assess how the derived parameters match other experimental observations (such as microscopy data). For example, the L-L fitting of wheat starches provided good S and fit uncertainty values, but made the invalid prediction of a long-tailing B-type granule peak, where a substantial proportion of B-type granules are within the 12-20 μm size range. Given that such giant B-type granules were never observed using microscopy, there is a strong biological basis to reject this fitting.

Our analysis of the *phs1* and *mrc* mutants provides an example where different samples within the same experiment optimally fit different fittings. The fact these fit different distributions compared to the wild type demonstrates that they have very different distributions. However, when making comparisons of quantitative parameters derived from the data, care should be taken if different fittings are employed between genotypes or treatments, since there could be different errors introduced by the different fittings. A better option in this example, since all fittings produced low S values, is to use the same fitting for all samples, but to state clearly that there could be error introduced by the curve fit being sub-optimal for some of the samples.

For rice and maize (Figure 7), the normal distribution fit better than the log-normal distribution, with low S scores. Although this suggests that these distributions could be mostly described with a normal distribution, there were some large granules that deviated from the distribution in rice. This could be a consequence of it forming compound granules (Matsushima et al. 2015). It is possible that the larger particles represent granules within the compound that have not fully dissociated from each other during starch extraction. It could also point to more complex mechanisms governing granule size. For example, physical interactions between the individual granules within the compound granules could influence the size of each particle, potentially causing a slight deviation from the theoretical distribution.

In the analysis of wheat starches, we were inspired by the work of Tanaka et al (2017), who developed a script for describing wheat starch with mixed distributions, with three log-normal distributions. Our script is different from this existing script in three ways. First, the Tanaka script was tested on data from a laser scattering instrument, whereas ours is tested on Coulter counter data. Second, our script strictly fits bimodal distributions, rather than trimodal distributions that include the small “C-type granule” peak, which was never observed in our experimental samples. It is possible that the occurrence of C-type granules is conditional (e.g., appears under specific environments). Finally, our script offers an L-N fitting, which we show provides the best fit for most Triticeae starches. Given these differences, it is up to the users to decide on which approach is most suitable for their purposes.

## 5 CONCLUSIONS

Our study provides a new method for the analysis of starch granule size distributions, and insights into best practices for quantitatively describing these distributions from different species. The script and relevant documentation are available online (see Methods). This resource will lead to more widespread, quantitative descriptions of granule size distributions within the research community.

## Acknowledgements

We are grateful to Martin Howard (John Innes Centre; JIC) and Richard Morris (JIC) for helpful discussions in preparing our script. We also thank JIC Horticultural Services for providing growth facilities and maintenance of plant material, the JIC Germplasm Resource Unit for providing seeds, and James Simmonds for assistance with growing the wheat samples. We also thank the National BioResource Project for barley, which is run by the Japanese Ministry of Education, Culture, Sports, Science and Technology (MEXT) for distributing the barley grains of Haruna Nijo cultivar. This work was funded through a John Innes Foundation (JIF) Chris J. Leaver Fellowship (to D.S), JIF Rotation Ph.D. studentships (to R.M and G.Y), a Biotechnology and Biological Sciences Research Council (BBSRC, UK) research grant BB/W015935/1 (to D.S.), and BBSRC Institute Strategic Programme grants BB/X01097X/1 and BB/X011003/1 (to the John Innes Centre).

## Supplemental Figures

**Figure S1:**
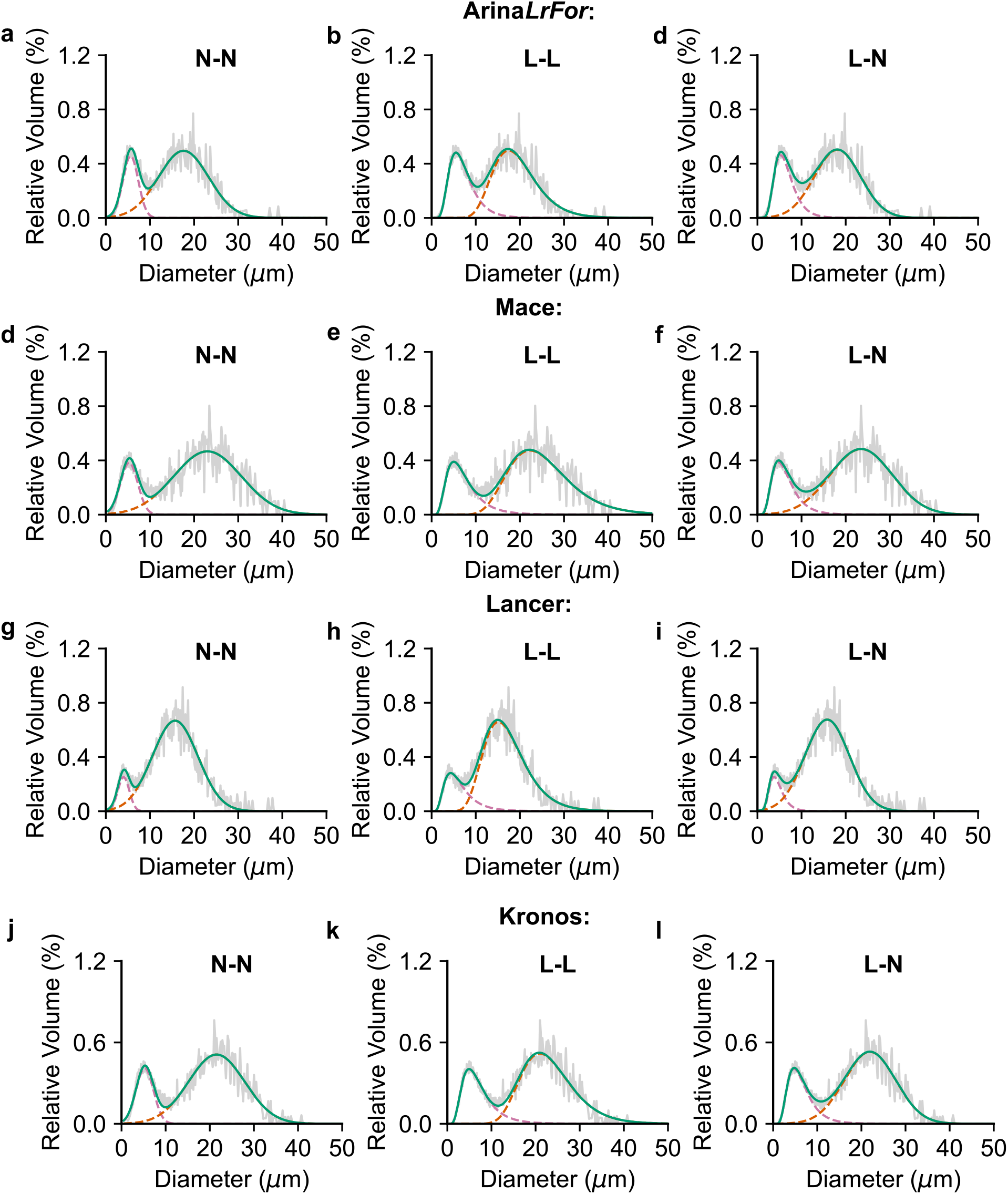
Three different bimodal curve fittings to a volumetric distribution of wheat starch. The normalised volumetric size distribution was determined using a Coulter counter on starch from wheat cultivars Arina*LrFor* (a-c), Mace (d-f), Lancer (g-i) and Kronos (j-l) (grey). **(a-l)** We fit two normal (a, d, g, j), two log-normal (b, e, h, k), or a log-normal and a normal (c, f, i, l) curves to the measured size distribution. The orange dashed curve represents the fitted A-type granule distribution, the pale pink dashed curve represents the fitted B-type granule distribution, and the green curve is the sum of these two distributions.

**Figure S2:**
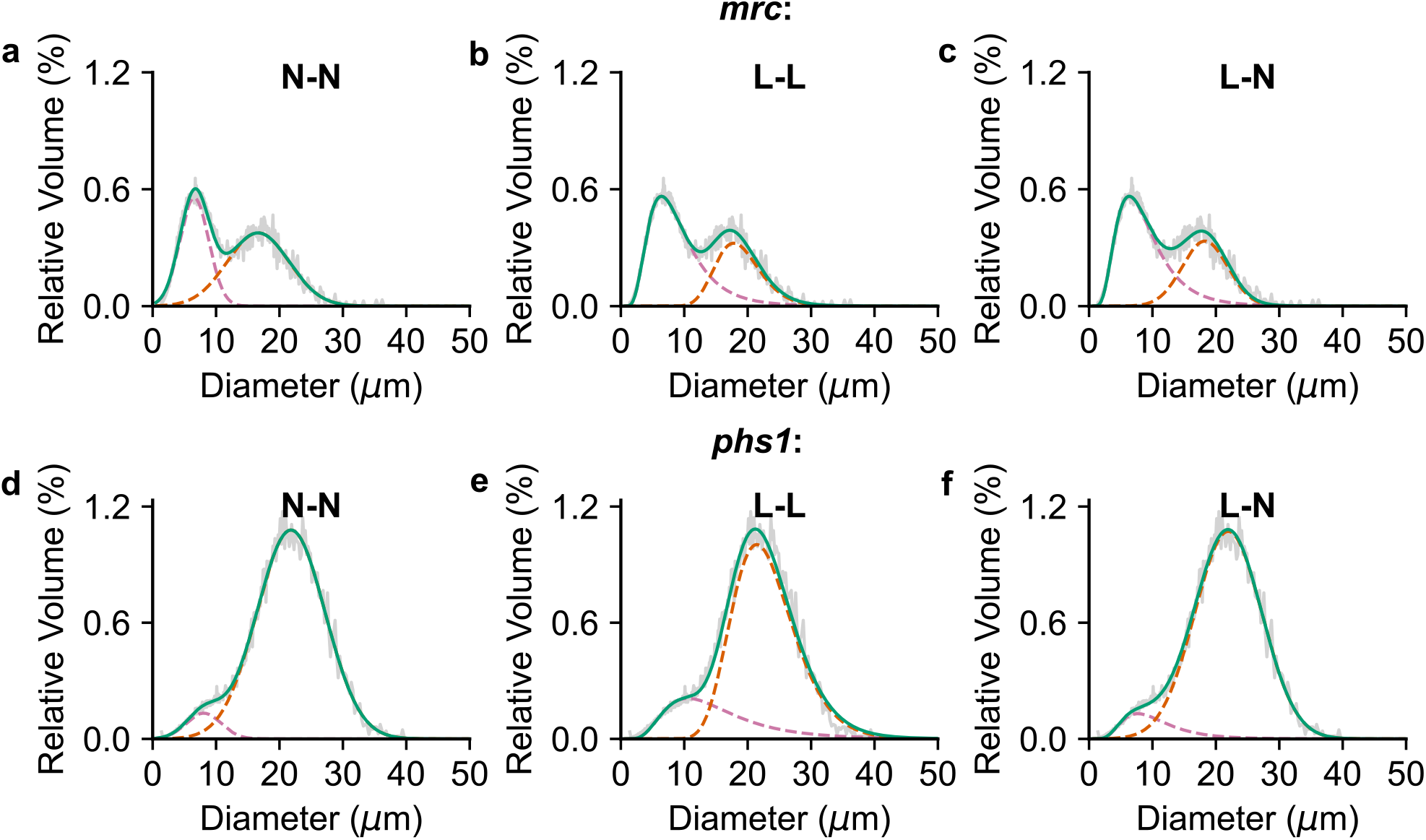
Three different bimodal curve fittings to a volumetric distribution of wheat mutants with altered B-type granule contents. The normalised volumetric size distribution was determined using a Coulter counter on starch from *mrc* (a-d), *phs1* (b-f) (grey). (grey). **(a-l)** We fit two normal (a, d), two log-normal (b, e), or a log-normal and a normal (c, f) curves to the measured size distribution. The orange dashed curve represents the fitted A-type granule distribution, the pale pink dashed curve represents the fitted B-type granule distribution, and the green curve is the sum of these two distributions.

**Figure S3:**
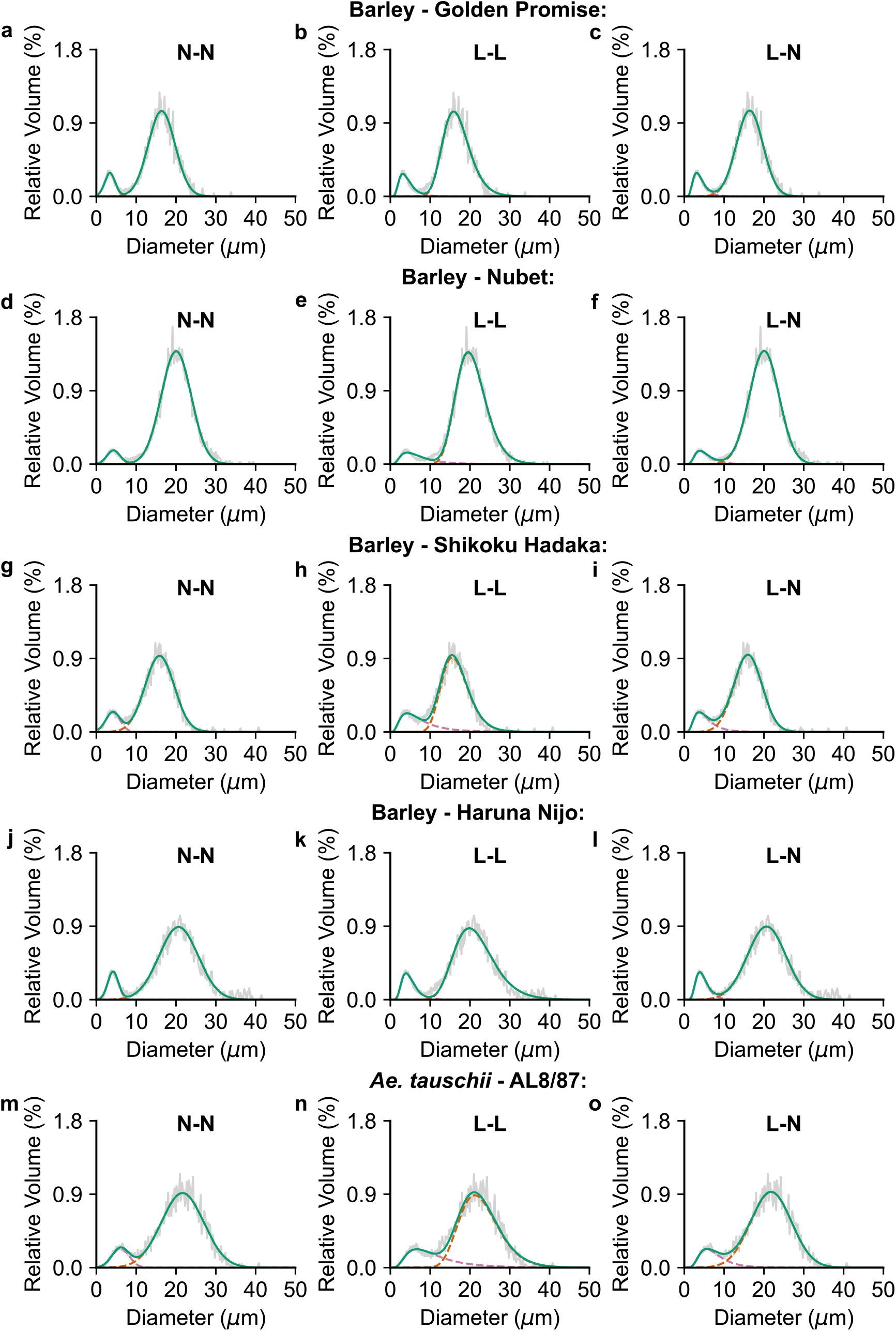
Three different bimodal curve fittings to a volumetric distribution of different Triticeae starches. The normalised volumetric size distribution was determined using a Coulter counter on starch from barley cultivars Golden Promise (a-c), Nubet (d-f), Haruna Nijo (g-i), Shikoku Hadaka (j-l), and the *Ae. tauschii* genome reference Al8/78 (m-o) (all grey). **(a-l)** We fit two normal (a, d, g, j), two log-normal (b, e, h, k), or a log-normal and a normal (c, f, i, l) curves to the measured size distribution. The orange dashed curve represents the fitted A-type granule distribution, the pale pink dashed curve represents the fitted B-type granule distribution, and the green curve is the sum of these two distributions.

